# The evolutionary origin of host association and polycistronic transcription in trypanosomatids

**DOI:** 10.1101/2025.10.27.684775

**Authors:** Saurav Mallik, Meir Sylman, Dvir Dahary, Orna Dahan, Shulamit Michaeli, Gerald F. Späth, Yitzhak Pilpel

## Abstract

Trypanosomatids (Kinetoplastids) encompass multiple lineages of parasitic protists with monoxenous or dixenous life cycles, infecting insects, vertebrates, and plants; in vertebrate hosts, some are intracellular, while others are extracellular. To understand the origin and diversification of their host associations, we integrated comparative genomics across 47 genomes. Results highlight that monoxenous, extracellular trypanosomatids originated from predatory ancestors through reductive evolution, which diminished their metabolic and hunting capabilities. Intracellularity and dixenous lifestyle convergently originated three times independently. Progressive consolidation of genes into polycistronic transcription units (PTUs) was a central innovation that began in early Glycomonada and expanded through chromosome fission-fusion and gene relocation/inversion. In present-day PTUs, protein complex subunits and metabolic pathway enzymes are positioned for co-expression in temporal synchrony, and chromosomes minimize colinear PTUs to counter transcriptional readthrough. Together, these results provide a time-resolved origin of host-association and polycistronic transcription in trypanosomatids, possibly through an intermediate phase of facultative parasitism.

## INTRODUCTION

Symbiotic associations between two or more different organisms have arisen many times independently across the Tree of Life and continue to shape biodiversity, ecosystem stability, and evolution of countless species^1–4^. These relationships encompass a wide range of diversity. Regarding the balance of benefits and costs, they range from being mutualistic to commensalistic to parasitic; and from facultative to obligate, depending on whether either partner can survive independently. The symbiont may be external or internal to the host (exo-/endosymbiosis), and endosymbionts can be extra- or intracellular. While a plethora of such host associations have been characterized, how they originate in evolution, and how one type of association transitions into another, remains poorly understood.

To investigate how host association arises and transforms over evolutionary time, we studied trypanosomatids, a group of flagellated protists of the class Kinetoplastea. They are obligate parasites displaying complex heteroxenous life cycles, typically involving a blood-sucking insect vector and a vertebrate/plant host with transmission through bites^5^. Within the insect guts, they exist in flagellated forms. Within the host, they are found either extra or intracellularly. The vector and host species, and the extra/intracellular location within the host, vary between specific trypanosomatid species. For example, *Trypanosoma brucei* is transmitted through tsetse flies and lives extracellularly in the mammalian blood and tissues. *T. cruzi* is transmitted through the feces of triatomine bugs and lives intracellularly within the mammalian host. *Leishmania spp* are transmitted through sand flies and exist intracellularly within mammalian macrophages. They are responsible for African and American trypanosomiasis^5^ and Leishmaniasis^6^ that, annually, result in ∼1.5 million new cases worldwide and ∼35,000 fatalities^7^.

The ecological and life-cycle features of these protists are coupled with their unusual genome organization and gene-expression strategies. They feature densely packed, largely intronless genes arranged in directional clusters, known as polycistronic transcription units, or PTUs, that resemble bacterial operons^8,9^. A PTU could harbor between a few to nearly a hundred genes that share a single transcription initiation and termination site and are transcribed into a polycistronic pre-mRNA, which is co-transcriptionally processed into mature, monocistronic mRNAs via trans-splicing and polyadenylation^10^. Consequently, trypanosomatids are thought to lack transcriptional regulation at the level of individual genes^11^. Their obligate host dependence is rooted in energy metabolism adapted to host-derived carbohydrates and limited intrinsic capacity to synthesize their own nucleotides, essential amino acids, lipids, vitamins, and cofactors^12–16^. Despite these insights into present-day trypanosomatid biology and host dependence, it remains unclear what their last free-living ancestors looked like, how host association first arose, and how their unconventional polycistronic transcription originated.

Here, we describe a systematic comparative genomics analysis suggesting that these parasitic trypanosomatids with polycistronic transcription originated from free-living predatory protists that featured a conventional, monocistronic eukaryotic transcription system. An ancestral genome-content reconstruction analysis illuminated the stepwise emergence and evolution of host association, the origin and expansion of polycistronic gene organization coupled with intron loss, and molecular underpinnings of gene organization within PTUs and PTU organization across chromosomes. Together, these results reveal the entwined evolution of trypanosomatids’ obligate host association and unconventional polycistronic transcription, advancing a systems-level view of the origins of parasitism.

## RESULTS

### A complex evolutionary history of host association in the trypanosomatid lineage

To illuminate the evolutionary diversification of trypanosomatids and their host associations, we conducted a systematic comparative genomics analysis. We obtained annotated proteomes of parasitic trypanosomatids and their closely related non-parasitic protists spanning Discoba and Fornicata clades and constructed a phylogenetic tree that represents species divergence (***Methods***). We screened publicly available genomic repositories^17–19^ and compiled annotated proteomes for 37 trypanosomatids and 10 related predatory or symbiotic protists (**Table S1**). We compared 503,423 proteins encoded in these genomes (each corresponding to a gene) using all-versus-all BLAST^20^ and clustered the resulting sequence-similarity graph into 32,894 protein families^21^ (**Data S1**). We used a maximum-likelihood method^22^ to reconstruct the species phylogeny from a concatenated alignment of 110 single-gene-copy families conserved in these 47 genomes (**Figure 1**). We rooted the species tree using the predatory protist clade Fornicata, which was previously reported as an outgroup to Discoba^23,24^. We used the TimeTree database^25^, which synthesizes data on species divergence times from thousands of published molecular-dating studies, to date the internal nodes of the species tree.

**Figure 1.**
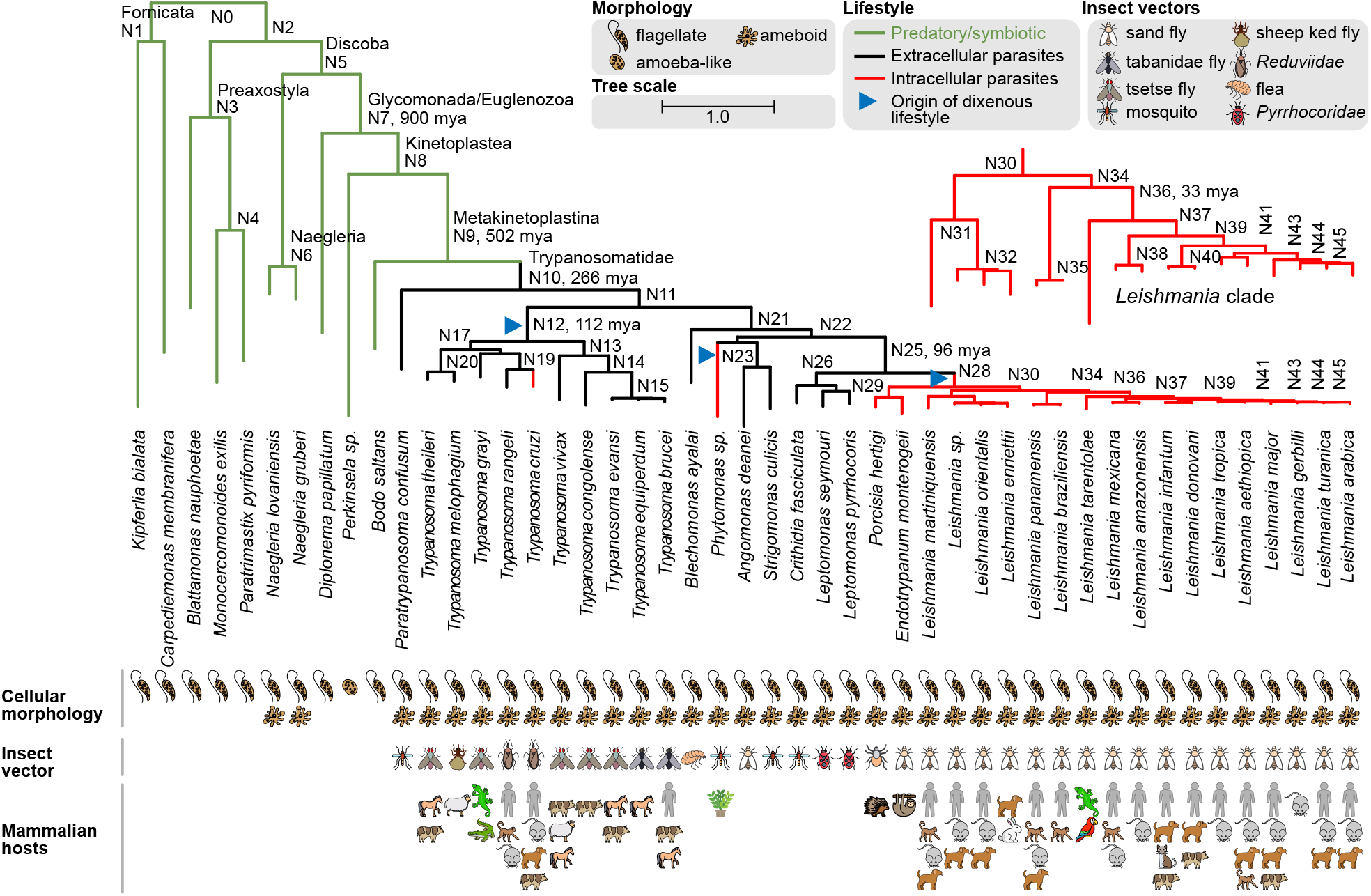
Evolutionary origin of trypanosomatids from non-parasitic ancestors. A species tree representing the evolutionary origin and divergence of Tryaonsomatids. It includes 47 protists with fully sequenced genomes (including 27 trypanosomatids and 10 predatory/symbiotic taxa). The tree was inferred using maximum-likelihood methods from a concatenated alignment of 110 single-copy protein families present in these genomes (***Methods***). Branch lengths represent the number of substitutions per 100 amino acids. All internal nodes have ≥80% bootstrap support. When relevant, NCBI taxonomy^40^-annotated clade names are provided alongside internal node numbers. When available, TimeTree database^25^-annotated origin dates are provided for the internal nodes (mya: million years ago). For each organism, known cellular morphologies, insect vectors, and mammalian/plant hosts are shown (sketched by the authors). For monoxenous parasites, only the insect vectors are shown. Branches representing intracellular and extracellular parasites are in red and black, whereas those representing non-parasitic protists (predatory/symbiotic) are in green. Blue triangles mark parsimoniously reconstructed origins of the dixenous lifestyle. For clarity, a zoomed-in *Leishmania* tree is provided.

Our species tree is consistent with recent large-scale reconstructions of protist and broader eukaryotic phylogeny^23,24,26,27^ and overcomes the limitations of previous attempts to decipher Kinetoplastea diversity using single-gene phylogenies, since any single gene is a poor proxy for species divergence, or alignment-free phylogenetic methods, which are less accurate^28–31^. It supports a monophyletic origin of parasitic trypanosomatids from predatory ancestors, whose descendants today occupy marine, terrestrial, and freshwater environments^32,33^. We found Bodonids, a bacterial predator in terrestrial/freshwater ecology, to be the closest free-living relative of trypanosomatids, as was previously reported^34^. Under a parsimonious reconstruction, obligate parasitism appears to have arisen once in the trypanosomatid common ancestor (TryCA; node N10), inferred as an extracellular, monoxenous parasite of insects. TimeTree^25^ places TryCA at ∼266 million years ago (mya), which is after insect origins (∼400 mya)^35^ but before mammalian origins (∼225 mya)^36^. These are consistent with the insect-early hypothesis^37^, which posits that TryCA was monoxenous (single-host), with dixenous parasitism (life cycles involving an insect vector and a vertebrate/plant host) evolving later. We infer three independent origins of a dixenous lifestyle within trypanosomatids (nodes N12, N28, and *Phytomonas*), at least ∼100 million years after TryCA. Obligate intracellular parasitism also arose three times independently (*T. cruzi, Phytomonas*, and N28). Extant descendants of earlier nodes, *i*.*e*., the last common ancestors of Discoba, Euglenozoa, and Kinetoplastea (N9-N5), span predators, endosymbionts, and parasites^38,39^, suggesting that ancestors of TryCA may have been free-living or facultative symbionts/parasites.

Taken together, these results highlight a monophyletic origin of parasitic trypanosomatids, followed by a gradual sophistication of their host association. The earliest trypanosomatids arose as monoxenous, extracellular insect parasites, and dixenous life cycles and intracellular invasion convergently evolved multiple times independently thereafter. These analyses provide a time-resolved origin history that is consistent with the insect-early hypothesis.

### Obligatory host association arose through episodic sparsening of core metabolic pathways and protein complexes

What were the molecular determinants of host association in trypanosomatids? To address this question, we compared per-taxon gene repertoires across the species tree to identify signatures of reductive evolution. In our dataset, the average gene count per genome decreased from 21,219 (±9,312) in the free-living outgroup protists to 10,124 (±1,309) in *Trypanosoma* and 8,638 (±214) in *Leishmania*. To reveal the identities of lost, retained, or potentially gained genes, leveraging protein-family annotations (**Data S1**), we inferred each family’s presence/absence and copy number across the internal nodes of our species tree using a maximum-likelihood approach^41^ (**Methods, Data S2**). This enabled two complementary views of gene loss: (i) loss of redundancy (the family is retained but shrinks in copy number via paralog loss; **Figure 2A**, blue heatmap) and (ii) loss of function (the family is entirely lost; **Figure 2A**, green bars). To relate these losses to the functional capacity in predatory ancestors, we classified the protist proteomes using KEGG^42^ (**Methods**).

**Figure 2.**
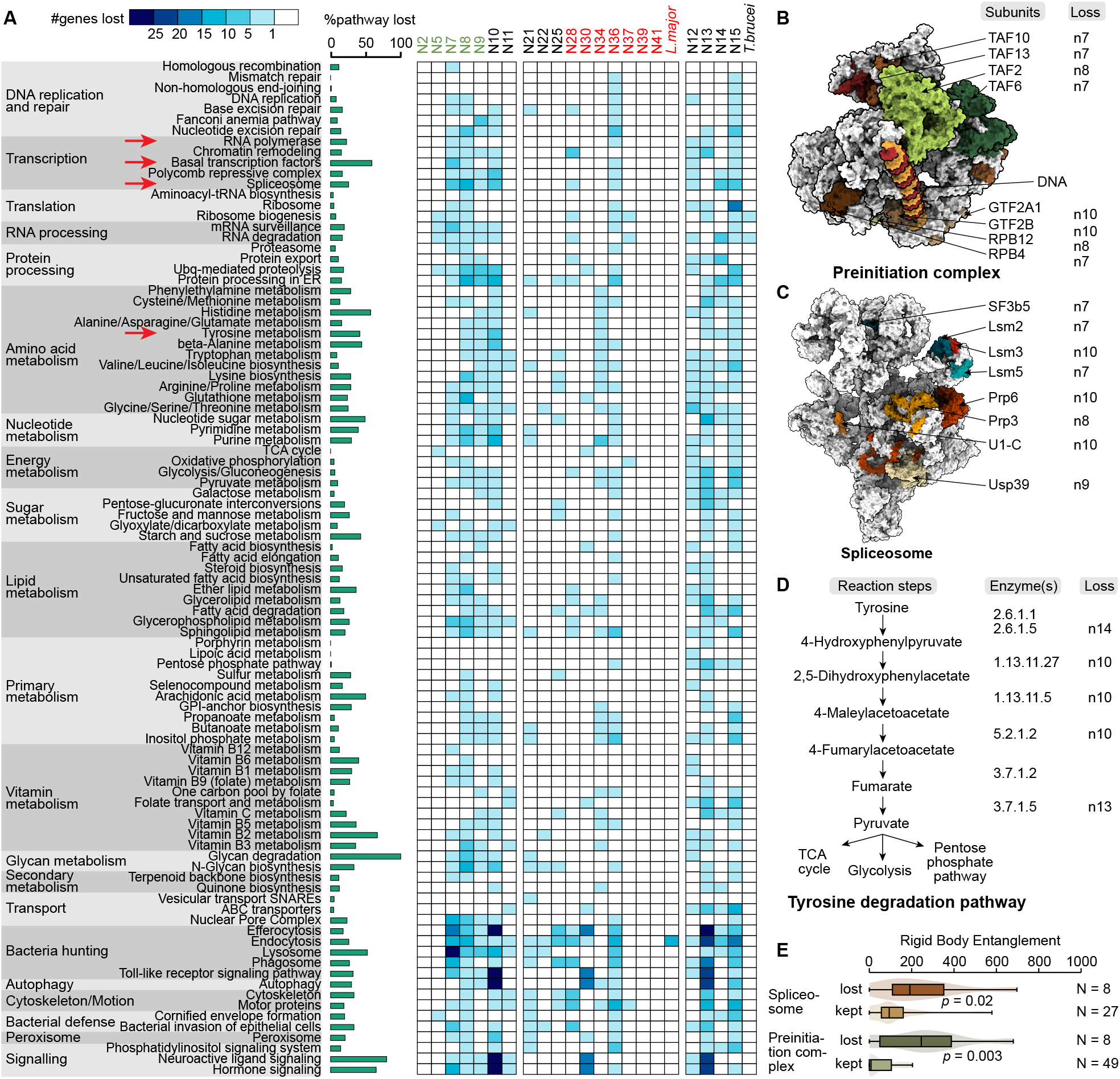
Origin of host association in trypanosomatids by reductive evolution. (**A**) A heatmap represents pathway loss in Trypanosomatid lineages leading to *L. major* and *T. brucei* due to reductive evolution. The core cellular pathways of free-living, heterotrophic protist *Naegleria gruberi*, representing the bulk of its gene repertoire, are highlighted on the left. These pathways were selected from the KEGG pathways collection^42^. The color in each cell depicts the number of gene loss events within each KEGG category, at various internal nodes of the species tree depicted in **Figure 1**. The green bars represent the % of each pathway lost in *L. major*, represented as the % protein families that were lost. (**B-D**) Reductive evolution of the transcription pre-initiation complex (PDB code: 7EDX^43^) (**B**), and Spliceosome (PDB code: 6QX9^44^) (**C**), and the Tyrosine degradation pathway (**D**) in trypanosomatids. These three systems are highlighted in panel **A** with red arrows. For the two complexes, conserved subunits are in grey, and those in color are lost in Trypanosomatid lineages leading to *L. major* and *T. brucei* due to reductive evolution. For each subunit in panels **B** and **C**, and each enzyme in panel **D**, the node of loss is mentioned. (**E**) For the lost and retained subunits of the preinitiation complex and spliceosome, violin plots compare their rigid body entanglements. This parameter recapitulates the number of ways a subunit can dissociate from the complex as a rigid body without causing steric clashes.

Results indicate an episodic, rather than gradual, decay of core cellular functions in trypanosomatids, including sparsening of metabolic pathways and simplification of macromolecular complexes (**Figure 2A**). The extent of sparsening varies across pathways. In core metabolism, trypanosomatids have retained energy metabolism and central biosynthetic pathways (glycolysis, the TCA cycle, oxidative phosphorylation, the pentose phosphate pathway, and fatty-acid biosynthesis), but lost major components of purine and pyrimidine metabolism, amino-acid biosynthesis, and vitamin/cofactor metabolism. Prior studies attributed their obligate host association to limited intrinsic capacity to synthesize nucleotides, amino acids, and vitamins, which are salvaged from the vector and host^12–16^. Trypanosomatids have also lost substantial portions of predation pathway genes (efferocytosis, endocytosis, lysosome, phagocytosis, and glycan degradation)^45,46^ that persist in predatory outgroup genomes.

When were these genes lost? Our analysis points to three major episodes of reductive evolution: N8-N10 (spanning ∼600 million years, by TimeTree^25^ annotations, preceding the obligate monoxenous, extracellular parasitism inferred at N10), N28-N36 in the *Leishmania* lineage (spanning ∼90 million years after obligate dixenous, intracellular parasitism at N28), and N12-N15 in the *Trypanosoma* lineage (spanning ∼100 million years after obligate dixenous, extracellular parasitism at N12). Notably, the N8-N10 episode features thousands of gene losses preceding obligate host dependence. One way to reconcile this is to speculate that a facultative symbiotic/parasitic relationship existed between N5 and N9, with sustained gene loss events culminating in obligate parasitism at N10. This hypothesis is also consistent with our earlier observation that descendants of N5-N9 include both free-living and parasitic taxa.

Our analysis also reveals compositional simplification of macromolecular assemblies. In particular, the transcription preinitiation complex (includes RNA polymerase II and the basal transcription factors, **Figure 2B**) and the spliceosome (**Figure 2C**) have lost multiple subunits in the trypanosomatid lineage. To test for structural/biophysical principles of subunit loss, we mapped lost components onto orthologous human complexes. To examine whether any structural/biophysical principles underlie loss versus conservation, for each subunit, we computed the number of ways in which it can dissociate from the rest of the complex as a rigid body without causing steric clashes^47^ (**Methods**). Higher or lower numerical values of this parameter, rigid body entanglement, reflect structurally peripheral or central subunits, and our results show that subunits that are lost today used to reside at the structural periphery (**Figure 2E**). By contrast, we did not detect an analogous principle for metabolic pathway sparsening. For example, in tyrosine degradation, the order of enzyme loss does not follow any simple rule. Enzymes catalyzing the first and last steps were lost in Trypanosoma spp after losses at intermediate steps (**Figure 2D**). In *Leishmania*, genetic knockout of the first enzyme tyrosine aminotransferase (EC 2.6.1.5) disrupts redox homeostasis and causes growth arrest^48^.

Taken together, obligate host association in trypanosomatids emerged through at least three episodes of reductive evolution, two of which post-date the origins of obligate dixenous parasitism in the *Leishmania* and *Trypanosoma* clades. Reductive evolution sparsened core metabolism, simplified protein complexes, and diminished predatory capabilities, contributing to a transition toward obligate host dependence.

### Origin of parasitic pathways in Trypanosomatid evolution

Trypanosomatids include several disease-causing parasites. *L. major*, for example, invades macrophages and transforms their phagolysosomes (vesicles that degrade engulfed microorganisms) into a parasitophorous vacuole, where they reside, acquire nutrients, hijack immune response pathways, and replicate^6^. When during trypanosomatid evolution did the corresponding pathogenicity factors originate? To address this, we examined the presence/absence and gene-copy numbers of protein families across the species tree (**Methods, Data S2**).

Results show that most protein families representing present-day pathogenicity factors were already present in the gene repertoire at ancestral nodes N0-N2, long before obligate parasitism arose at N10 or the proposed facultative parasitism arose at N5 (**Figure 3**). These include numerous surface and secreted factors: lipophosphoglycans (prevent phagosome maturation and acidification^49^), the major surface antigen leishmanolysin (promotes macrophage invasion and immune evasion^50^), and glycoinositolphospholipids (dampen macrophage activation^51^). The same applies to key metabolic and stress-response components: arginase (depletes L-arginine to reduce microbicidal nitric-oxide biogenesis^52^), superoxide dismutases (counter the host oxidative burst^53^), peroxiredoxins (detoxify peroxides^54^), and trypanothione reductase (supports detoxification of reactive oxygen species^55^), as well as modulators of host signaling, including cysteine/serine/aspartyl proteases (degrade host MHC class II molecules^56^ and phosphatases (dephosphorylate host signaling proteins^57^).

**Figure 3.**
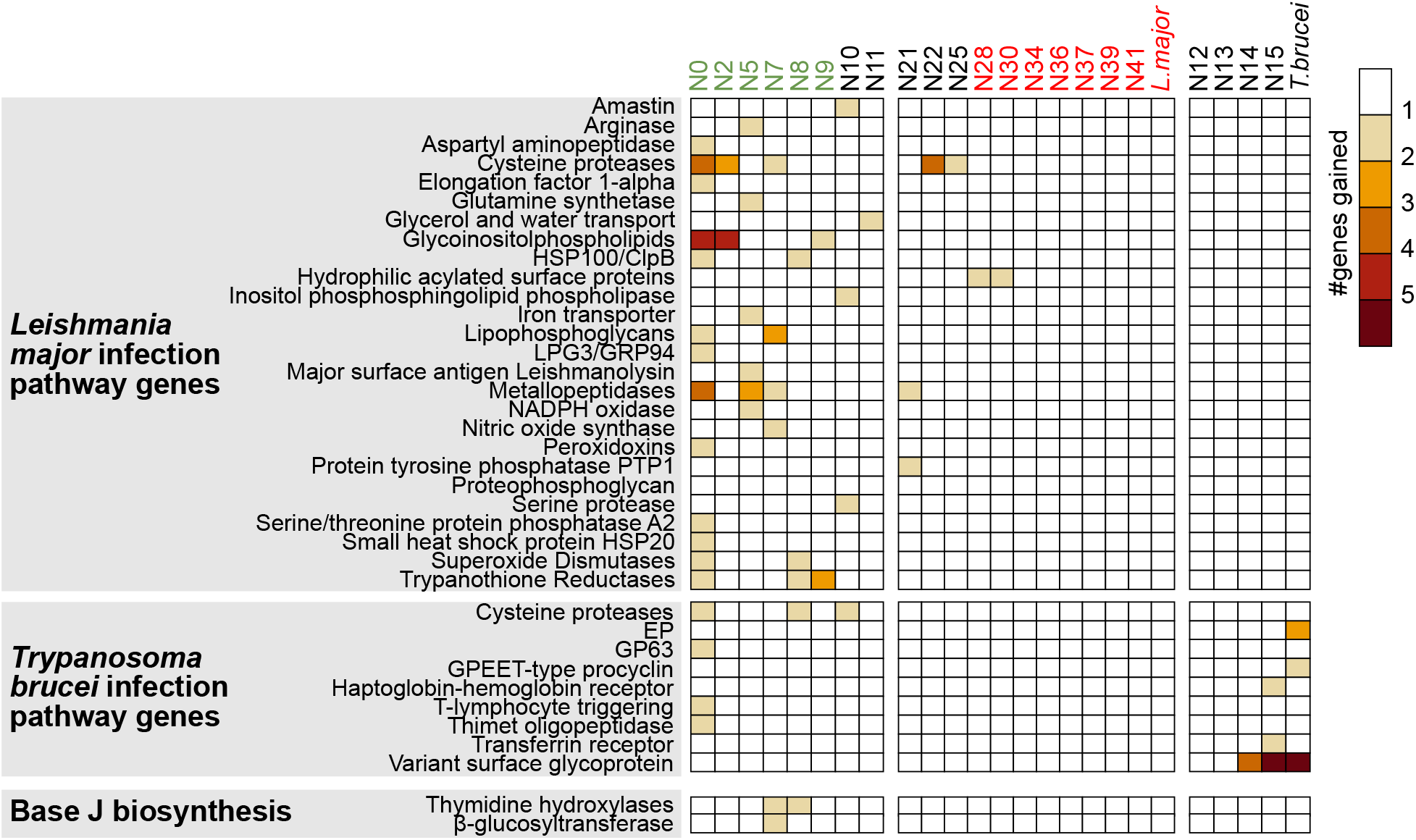
Origin of pathogenicity factor genes in trypanosomatids. A heatmap represents gene gain events in Trypanosomatid lineages leading to *L. major* and *T. brucei* in pathways related to their infection pathway. The color in each cell depicts the number of gene gain events within each category, at various internal nodes of the Trypanosomatid species tree (**Figure 1**).

By contrast, some hallmark pathogenicity factors of the extracellular parasite *Trypanosoma brucei* appear relatively recently (**Figure 3**). These include the dedicated surface proteins EP procyclin (specific to the procyclic form in the tsetse fly^58^ and variant surface glycoproteins (VSGs), which form a dense bloodstream-form coat enabling immune evasion via rapid antigenic variation by switching among VSG genes^59^.

The prevalence of pathogenicity-factor genes before the emergence of parasitism itself supports a co-option model: in free-living predatory ancestors, these proteins plausibly mediated defense against capture and digestion. As host association evolved, the same activities may have been co-opted to subvert macrophage phagolysosomes and blunt immune response, thus converting predator-evasion tools into parasitic effectors.

### Genome Tramlining in Trypanosomatids

A key feature of trypanosomatid genomes is their polycistronic transcription system^8,60^. Nearly 99% of all *L. major* genes occur in PTUs^8^ (**Figure 4A**). How did such an unusual genome architecture originate in evolution? To address this question, we systematically compared gene orientations across our species tree.

**Figure 4.**
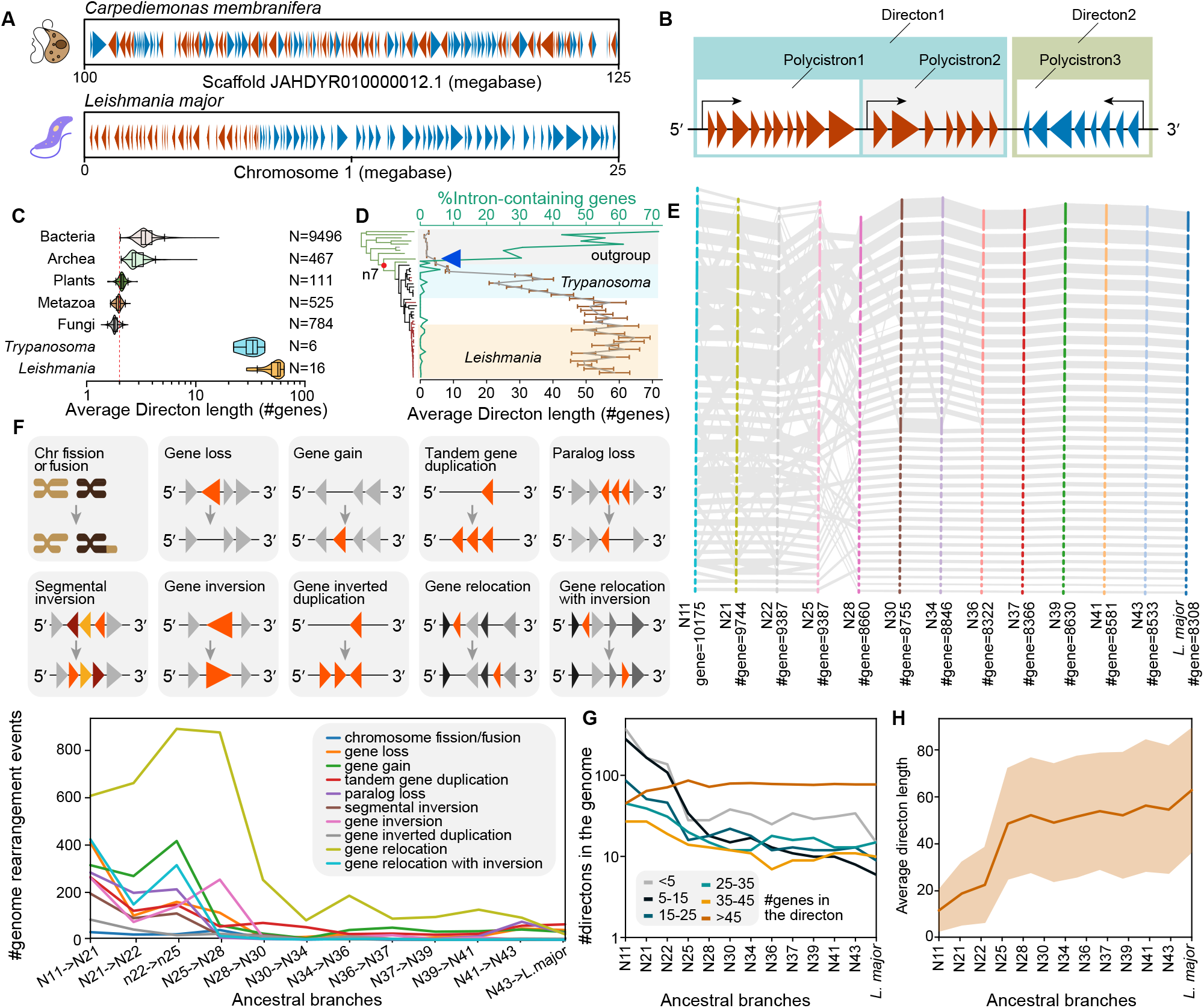
Incremental evolution of genomic tramlining in Trypanosomatid lineage. (**A**) For the heterotrophic protist *C. membranifera* and obligatory parasite *L. major*, the red or blue triangles represent annotated genes and their orientations within a 25 megabase region of their respective genomes. (**B**) A schematic representation of a directon, which includes one or more co-linear PTUs. (**C**) Violin plots represent the distribution of average #genes per directon across the three domains of life. (**D**) The grey line and the orange bars represent the average #genes per directon and the corresponding standard deviations in the 47 genomes included in our species tree. The green line represents the %intron containing genes in the genome. The blue arrow marks the evolutionary onset of genome tramlining. (**E**) The evolution of chromosome numbers and synteny in the Trypanosomatid lineage leading to *L. major*. The rightmost stack of blue vertical bars represents the 36 chromosomes in the *L. major* genome, which includes a total of 8303 protein-coding genes. The remaining stacks represent the same for phylogenetically reconstructed ancestral genomes (N11, N21, … N43, see ***Methods***). Synteny blocks are shown in grey. (**F**) Top, schematic diagrams represent different types of genome rearrangement events that can contribute to genome tramlining in trypanosomatids. Except for chromosome fission/fusion, the remaining events are drawn such that the gene(s) involved (not involved) in the event are in orange (gray). Bottom lines represent the frequency of different phylogenetically reconstructed genome rearrangement events occurring at the ancestral branches of the Trypanosomatid species tree. (**G**) Lines represent the frequency of directons of different length ranges in the extant and reconstructed genomes across the Trypanosomatid species tree. (**H**) The line and the surrounding shaded area represent the average #genes per directon and the corresponding standard deviations at the ancestral nodes of the species tree.

The transcription start and stop sites in Trypanosomatid genomes (*i*.*e*., PTU boundaries) are designated by chromatin marks^61^ and modified DNA base J^62^, which remain uncharacterized for most protists in our tree. To overcome this, we introduced a proxy: a *directon*, defined as a group of consecutive, co-linear genes on the same DNA strand (**Figure 4B**). A directon can comprise one or more co-linear PTUs. Examining 11,583 genomes across all three domains of life, we found that the average #genes/directon is ∼2 in multicellular eukaryotes, which reflects near-random strand assignment of genes, and ∼3-4 in prokaryotes, which reflects operons. Trypanosomatids stand out, reaching ∼33 and ∼56 genes/directon in *Trypanosoma* and *Leishmania* (**Figure 4C**). This contrast is recapitulated within our species tree (∼2 genes/directon in outgroup predatory protists; **Figure 4D**). These results highlight that both the fraction of polycistronic genes and the average #genes/PTU increased along the trypanosomatid lineage. We term this evolutionary process as *genome tramlining*.

When did the first PTUs originate and genome tramlining begin? To address this, we traced the genetic origin of enzymes in the base J biosynthesis pathway. Base J is synthesized in two enzymatic steps^63^. First, thymidine hydroxylases oxidize methyl groups on specific thymines, producing hydroxymethyl-deoxyuridine (HOMedU). Next, a β-glucosyltransferase converts HOMedU into base J by adding a glucose molecule. Our phylogenetic analysis traced the origin of both enzymes to Glycomonada (N7, **Figure 3, Data S2**). An initially acquired single-copy thymidine-hydroxylase gene later duplicated in Kinetoplastida (N8), yielding the JBP1/2 paralogs currently found in the *L. major* genome. We also found that the expansion of polycistronic architectures was associated with a drastic loss of introns. Free-living descendants of N0-N5 feature conventional monocistronic genomes (∼2 genes per directon) and ∼53% intron-containing genes per genome. By contrast, Kinetoplastida and diplonemids, descendants of N7, harbor tramlined genomes (≥7 genes per directon) and 34% and <1% intron-containing genes. (**Figure 4D**). These observations suggest that the first PTUs originated and genome tramlining began in Glycomonada (N7), ∼600 million years before the origin of obligate parasitism at N10. A potential explanation for the parallel decline in *cis* splicing and increase in *trans* splicing (necessitated by tramlining) is that if both *forms of* splicing occur for the same gene, they compete for the same splice sites, risking the loss of exons.

Which genetic mechanisms facilitated genome tramlining, and what were their relative contributions? To address this question, we reconstructed the ancestral gene orientations across our species tree (***Methods***). Leveraging gene orientations in extant trypanosomatid genomes and our protein-family reconstructions (**Data S1**), we used a maximum-likelihood method^41^ to infer consecutive gene pairs and their orientations at each bifurcation. We then reintegrated these pairs into full chromosomes (**Data S3**) following the AGORA pipeline^64^. Our reconstructions are limited to N11 because chromosome-level genome assemblies are unavailable for deeper nodes.

We found that chromosome fission/fusion events dominated early trypanosomatid evolution (nodes N11-N28); these events largely subsided at N30, after which descendant *Leishmania* species maintained high gene synteny (**Figure 4E**). Using chromosome-level ancestral reconstructions, we inferred the specific sequence of rearrangements that produced extant *Leishmania* and *Trypanosoma* genomes (**Figure 4F, Methods**). We found that relocation (with or without inversion), inversion, loss, gain, or duplication of genes occurred in a sequence that progressively tramlined the genomes in the lineages leading to *L. major* and *T. brucei*. The count of short directons (≤45 genes) gradually decreased in their genomes, whereas long directons (>45 genes) increased (**Figure 4G**). The average #genes/directon rose from ∼12 at N11 to ∼55 in *Leishmania* (**Figure 4H**). Results for *T. brucei* and *T. cruzi* lineages are shown in **Figure S1**. These results suggest that genome tramlining proceeded via the gradual fusion of smaller directons into increasingly longer ones.

Taken together, these results uncover the fascinating evolutionary history of genome tramlining in trypanosomatids. Formation of the first polycistrons coincides with the genetic origin of base-J biosynthesis enzymes in early Glycomonada, long preceding the rise of obligate parasitism. A variety of genetic events facilitated the expansion of PTUs in parasitic lineages by adding more genes to existing units, primarily through gene relocation and inversion, to the point that in the *L. major* genome, more than 99% of protein-coding genes have lost monocistronic identity.

### The molecular principles of present-day Trypanosomatid genome architecture

What were the molecular principles of genome tramlining, *i*.*e*., in what order are individual genes organized within PTUs, and individual PTUs arranged along chromosomes? Prokaryotic operons typically comprise 3-4 genes (**Figure 4C**). Given an average prokaryotic gene length of ∼800 nt^65^ and transcription elongation speed of 20-90 nt/s^66^, genes within the same operon are transcribed with an approximate delay of ∼1-2 minutes, which is shorter than the median bacterial mRNA half-life (∼3-4 min)^67,68^. In other words, genes within an operon are produced at balanced stoichiometry and, physiologically, in near temporal synchrony. Accordingly, operons tend to harbor functionally related genes, such as subunits of the same complex or enzymes in the same pathway^69^.

By contrast, trypanosomatid PTUs do not follow this rule. Using eukaryotic elongation rates of 50-90 nt/s^70,71^, transcribing an average PTU in *L. major* is estimated to take ∼38 minutes (**Figure S2A**), which exceeds the median mRNA half-life (∼16 min)^72,73^. Thus, genes within a PTU are produced at similar stoichiometry but, physiologically, not necessarily at the same time, except for consecutive genes. This suggests that large complexes and pathways, which are systems requiring both stoichiometric balance and temporal synchrony, would need to be split across multiple PTUs, rationalizing why trypanosomatid PTUs appear to contain functionally unrelated genes^74^. To test this, we compiled enzyme/subunit compositions for annotated eukaryotic metabolic pathways^75^ and heteromeric complexes^76^ and mapped their orthologs in the *L. major* genome (***Methods***). We found that the number of subunits/enzymes in a complex/pathway positively correlates with the number of PTUs harboring their genes (**Figure S2B-C**). Moreover, gene indices of co-pathway/co-complex pairs tend to be significantly more adjacent than expected at random when they reside in the same PTU (**Figure 5D-E**), indicating a tendency to be consecutive. Notably, a similar correlation holds even when such pairs reside in different PTUs (**Figure 5D-E**), implying that these genes lie at comparable distances from their respective transcription start sites and may therefore be expressed in temporal synchrony. Could these trends arise by chance? We evaluated this with a permutation test that generated a null distribution of gene-index correlations. Specifically, in each of 1000 iterations, we sampled 1000 random gene pairs from the genome and recomputed the gene-index correlations for each iteration. The observed correlations (circle and star, **Figure 5F**) exceed the null expectations (violin plots, **Figure 5F**) with strong statistical support (right-tailed Z-test *p*-value < 1E−200 for each comparison), indicating that complex subunit and pathway enzyme arrangements did not arise by random placement.

**Figure 5.**
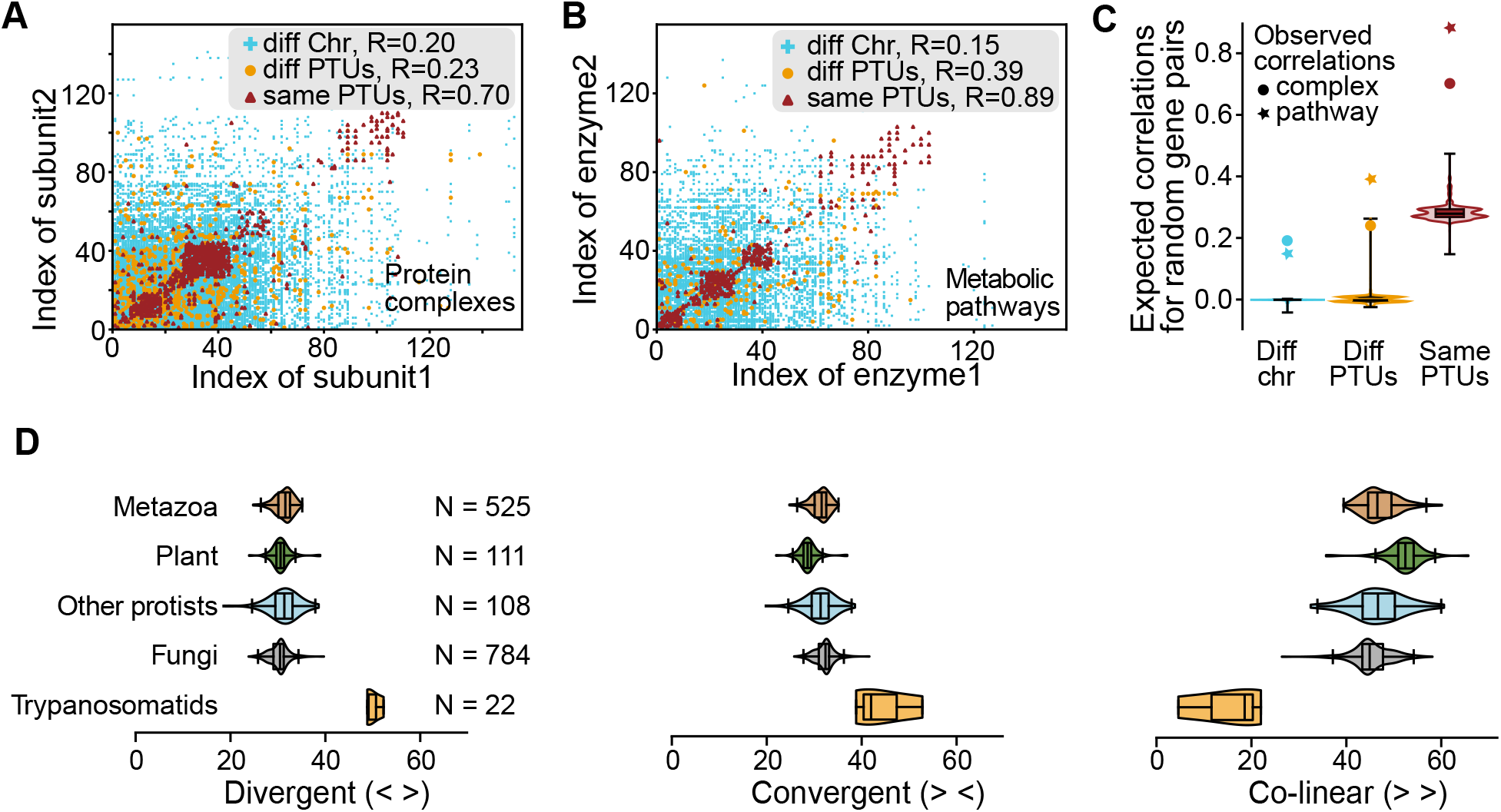
Gene organization in PTUs and PTU organization in chromosomes. (**A-B**) For subunit pairs of the same heteromeric complex (**A**) or enzyme pairs of the same metabolic pathway (**B**), a scatter plot shows their gene indices in their corresponding PTUs. Red, orange, and blue colors represent enzyme/subunit pairs encoded in the same PTU, different PTUs but the same chromosome, and different chromosomes, respectively. (**C**) These observed correlations (circle and star: co-complex subunit and co-pathway enzyme pairs) are compared to the null expectation. The latter were computed as the gene index correlations for 1000 randomly sampled gene pairs over 1000 iterations and are shown as violin plots. The boxes and whiskers span 25%-75% and 5%-95% of the respective distributions; the horizontal lines represent the medians. (**D**) The violin plots compare the % of divergent, convergent, and co-linear transcriptional units across eukaryotic genomes. For non-trypanosomatids, the transcriptional units are individual genes. Violin features are the same as panel **C**.

How are different PTUs organized along a chromosome? To address this, across the domain Eukaryota, we compared the orientations of consecutive transcriptional units. In typical eukaryotic genomes, individual genes are the transcriptional units, and their chromosomes contain ∼25% divergent, ∼25% convergent, and ∼50% colinear gene pairs (**Figure 5F-H**), a pattern consistent with near-random strand assignment. By contrast, trypanosomatid chromosomes are enriched for divergent and convergent PTUs (∼45% and ∼40%) and depleted for colinear PTUs (∼15%). We reasoned that a genome with naturally leaky transcription termination would be under selection to avoid colinear transcriptional units. Trypanosomatids are known to lack conventional eukaryotic terminators, and RNA polymerase II stop sites at PTU boundaries are instead epigenetically marked. Distinct polymerase leakage has indeed been observed in trypanosomatids, both naturally^77^, and upon base J disruption^62,77,78^. However, to what extent their leakage compares to eukaryotes with conventional terminators remains unknown. Additionally, divergent gene pairs in yeast were shown to be often co-degraded through promoter-mediated transcription-degradation coupling^79^. A similar mechanism may exist in trypanosomatids, which are heavily reliant on post-transcriptional control of mRNA levels.

Taken together, these patterns suggest that genome tramlining in trypanosomatids was governed by the principles of transcriptional control. Individual genes are arranged within PTUs to facilitate temporally synchronized co-expression of protein-complex subunits and pathway enzymes. PTUs are organized along chromosomes to minimize transcriptional readthrough.

## DISCUSSION

Trypanosomatids are a group of parasitic protists with “unusual biology”: they switch cellular morphologies within insect vectors and mammalian hosts, and feature polycistronic transcription combined with trans-splicing. Our systematic phylogenetic analyses show how these unusual features gradually evolved from non-parasitic ancestors featuring more conventional biology in these respects. These ancestors appear to have been free-living predators that preyed on bacteria or other protists, catalyzed their own energy metabolism, and synthesized necessary metabolites such as NADPH, pentose sugars, fatty acids, nucleotides, amino acids, vitamins, and cofactors. Their genomes also harbored genes for the synthesis of lipophosphoglycan, arginase, superoxide dismutase, peroxiredoxin, and trypanothione reductase genes, factors that their present-day descendants, *Leishmania*, use to resist digestion within macrophages. Taken together, this ancestral gene repertoire indicates marked phenotypic plasticity: these organisms could both prey on other microbes and defend themselves against being captured and digested.

Intriguingly, the evolutionary transition from these free-living protists to their obligate parasite descendants is characterized by a sustained period of reductive evolution that sparsified core metabolism, simplified protein complexes, and diminished predatory capacity. As this process continued, capabilities for nucleotide, amino-acid, vitamin, and cofactor biosynthesis were lost, contributing to a transition toward obligate host dependence. We propose that this reductive evolution was set in motion by an initial shift to facultative parasitism. Our results indicate that this shift was not driven by a sudden origin of new genes but by novel use of pre-existing genes, thus featuring phenotypic plasticity and co-option^80^. The transition from facultative to obligate parasitism may have occurred via genetic assimilation and/or the Baldwin effect^81–83^. In genetic assimilation, plastic phenotypes are fixed in the genome under persistent selection, while the Baldwin effect posits that plasticity enables temporary survival, buying time for heritable variants that support the lifestyle to arise and be selected. Taken together, our results highlight a possible phase of facultative parasitism as an evolutionary stepping-stone toward obligate parasitism in trypanosomatids, a trajectory observed in other parasitic clades as well^81,84,85^.

Our results further highlight that polycistronic transcription in trypanosomatids arose from ancestors with conventional monocistronic transcription, through a process we term genome tramlining: progressive loss of individual promoters and consolidation of genes into long, co-transcribed PTUs. We traced the molecular toolkit for this system to an early Glycomonada, when the base-J biosynthesis pathway first evolved. Base J epigenetically defines PTU boundaries today. Consistent with this, all extant Glycomonada descendants exhibit nearly intronless, tramlined genomes, albeit to very different degrees. This trait became exceptionally pronounced in parasitic trypanosomatids, where genomic rearrangements have organized the vast majority of intronless genes into large PTUs, erasing their monocistronic identities. By contrast, other Glycomonada descendants, free-living diplonemids and bodonids, and the intracellular endosymbiont *Perkinsela*, display milder tramlining and retain individual transcription of many genes. Notably, bodonids are primarily flagellated^86^, *Perkinsela* is only amoeboid^87^, and diplonemids switch from an amoeboid feeding/reproductive form to a flagellated foraging form upon starvation^86,88^. These observations are consistent with the hypothesis that extreme genome tramlining was specifically selected as an adaptation to a complex parasitic lifecycle requiring rapid morphological switching between insect and mammalian hosts. We propose that by eliminating the need to coordinate thousands of individual promoters, the parasites could trigger a global overhaul of their transcriptome and change morphology relatively quickly.

Finally, we show that organization of the tramlined genomes is governed by principles of transcriptional control at two levels. First, at the gene level, although individual PTUs contain functionally unrelated genes, subunits of the same protein complex or enzymes of the same metabolic pathway, while split across multiple PTUs, tend to be positioned at similar distances from their respective transcription start sites, either on the same or different PTUs. Such an intriguing gene arrangement indicates a fail-safe or a buffering mechanism, whereby fluctuation in the transcription rate of one PTU could be mirrored by inducing a similar fluctuation in the other, thus preserving the subunits’/enzymes’ stoichiometric balance as well as temporally synchronized expression. Indeed, PTU-level transcriptional fluctuations do occur in trypanosomatids^77,89,90^. Second, at the chromosome level, PTUs themselves are predominantly ordered in divergent and convergent arrangements while avoiding colinear arrangements. Prevalent divergent arrangements may refer to mechanisms akin to transcription-degradation coupling of divergent gene pairs observed in yeast^79^. Avoidance of colinear arrangements minimizes transcriptional readthrough when RNA polymerase II leaks from one PTU into the next. This architectural feature naturally implies a leaky transcription termination system. Indeed, distinct polymerase leakage has been observed in multiple trypanosomatids, both naturally^77^, and upon disruption of Base J biosynthesis^62,77,78^.

In summary, we have unveiled a complex evolutionary history of the origin and establishment of obligatory host dependency in trypanosomatids. Our framework demonstrates the power of integrating phylogenomics with systems biology and provides key insights into long-standing questions, including the stepwise origins of parasitism, the simplification of protein complexes in reductive evolution, and the mechanism and molecular principles of genome tramlining.

## Acknowledgments

We thank Peter J. Myler for his insight into the Base J biosynthesis pathway and all members of the Michaeli, Späth, and Pilpel lab for helpful discussions. This work was supported by the ERC 2022-SYG DECOLeishRN grant to S.M. (Bar-Ilan), G.F.S., and Y.P., and by the Minerva Center for Live Emulation of Evolution in the lab to Y.P.

## Author contributions

S.M. (TAU), M.S., D.D., O.D., and Y.P. conceived and conceptualized the research. S.M. performed the research, prepared the figures, and wrote the first draft. S.M. (Bar-Ilan), G.F.S., and Y.P. acquired funding; Y.P. supervised the study. All authors discussed and commented on the results and edited versions of the paper.

## Data and Materials availability

All data used or generated in this study are provided as Supplementary Materials.

