## Supplementary Methods for "The evolutionary origin of host association and polycistronic transcription in trypanosomatids"

**This Supplementary Materials file includes**

- Supplementary Data File Legends (SX-SY)
- Supplementary Methods
- Supplementary Table S1
- Supplementary Figures, SX-SY
- Supplementary References X-Y

**Supplementary Data File Legends**

**Data S1.** A list of the protein families identified across 47 protist genomes.

**Data S2.** Inferred gene copy numbers for protein families with KEGG functional annotations.

**Data S3.** Chromosome-scale reconstruction of ancestral genomes across the trypanosomatid species tree.

**Materials and Methods**

**Taxon sampling and species tree reconstruction**

We sampled 47 organisms with fully sequenced genomes (**Table S1**), following our previously published methodology^1,2^. Briefly, we sampled 37 representative proteomes for known parasitic trypanosomatids within the clade Kinetoplastea. We also sampled 10 representative proteomes of their non-parasitic protists from the clade Discoba and Fornicata. In these proteomes, each protein corresponds to a gene. BUSCO scores assessed the genome completeness of these proteomes. These scores were >95% for *Leishmania* and *Trypanosoma* taxa, but ∼80% for the non-parasitic genomes. Low BUSCO scores are typical for protists due to biological factors, such as gene loss in highly adapted or parasitic species, as well as technical limitations, including inadequate BUSCO datasets for specific protist groups and complications in orthology assessments resulting from their rapidly evolving genomes^3–5^.

To construct the species tree, we first identified protein families among these 47 genomes using the OrthoFinder package v2.5.5^6^ (parameter settings: -M msa -S mmseqs -A muscle -I 1.5). Briefly, OrthoFinder analyses include two steps. In the first step, an all-*versus*-all BLAST^7^ search compares all annotated protein sequences within these genomes^7^. We filtered the resulting BLASTp hits by removing all query-subject pairs for which the alignment coverage was <30% of either the total query or subject sequence length. This step is expected to significantly improve phylogenetic trees and orthology inference in the subsequent steps^8,9^. In the second step, based on these modified BLAST tables, we clustered the remaining sequences (using an inflation parameter of 1.5) into 30,773 protein families (**Data S1**).

Out of these 30,773, we extracted protein sequences of 110 single-copy families and subsequently aligned them using the tcoffee program^10^, which combined the outputs of PCMA^11^, MAFFT^12^, ClustalW^13^, POA^14^, Muscle^15^, and tcoffee^10^ alignment programs (parameter settings: -mode mcoffee -gapopen 20 -gapext 5). Each alignment was then trimmed using trimAL^16^ (parameter settings: -gt 0.6 -cons 50) to remove gap-majority columns. The resulting trimmed alignments were concatenated into a super-alignment, which was then finalized by removing the 20% most compositionally heterogeneous sites^17,18^. The final super-alignment included 93,707 columns. ModelFinder was then employed to find the best substitution model, which, based on Bayesian Information Criterion (BIC), turned out to be LG+F+I+G4 (-LnL = 2461441.24, BIC = 4924164.64). Maximum Likelihood phylogenetic inference was made using IQtree v1.6.12^19^ using 1000 ultrafast bootstraps (-m LG+F+I+G4 -bb 1000). The resulting tree was rooted using the two Fornicata species (*K. bialata* and *C. membranifera*).

**Ancestral gene repertoire inference and annotation**

We annotated the ancestral gene repertoire using a maximum-likelihood approach, which infers the gene copy numbers for the ancestral nodes of the species tree. We used the PastML package^20^, which, given a rooted phylogenetic tree with annotated leaves (gene copy-number annotation), infers ancestral copy numbers on the internal nodes using a character evolution model (ML method = Marginal Posterior Probabilities Approximation or MPPA, character evolution model = Estimate-from-Tips or EFT). Each inference is assigned a posterior probability (PP) score; pp ≥ 0.5 was used as the threshold to infer gene presence.

To infer the functions of these genes, we conducted KEGG pathway annotation for the free-living and parasitic proteomes using KEGG Mapper^21^ and BlastKOALA^22^. We integrated these mappings with proteome-scale KEGG^23^ annotations for *Leishmania major*, *Trypanosoma brucei*, and *Naegleria gruberi*.

**Annotating protein complexes and metabolic pathways**

We obtained subunit composition annotations for eukaryotic protein complexes from the Complex Portal database and mapped each subunit to the trypansomatid protein families by reciprocal BLAST^7^, using a ≤1E−10 e-value cutoff. From MetaCyc^24^, we similarly obtained enzyme gene annotations for all known eukaryotic metabolic pathways and similarly mapped them to the trypansomatid protein families.

**Comparing the average number of genes per directon across the tree of life**

Since PTU boundaries remain uncharacterized for most protists in our tree, we introduce an alternative metric: a directon, defined as a group of consecutive, co-linear genes on the same DNA strand. We obtained fully sequenced high-quality genomes of 9496 bacteria, 467 archaea, 111 plants, 525 metazoa, and 784 fungi from the Ensembl database^25^. We used an in-house Python script to identify directons residing in each chromosome. Finally, we computed the average number of genes per directon for each genome. The same analysis was repeated across 47 genomes included in the species tree.

**Intron analysis**

We collected intron annotations from the genome annotation files for each genome. Only ≥50 nt introns were considered in our analyses.

**Reconstructing ancestral chromosomes**

We inferred ancestral gene adjacencies with orientation using the PastML package^20^ and then assembled full ancestral chromosomes using the AGORA contiguity/linearization pipeline^26^. For 22 trypanosmatid genomes with chromosome-level assembly, we compiled gene orientations from the corresponding genome annotation files and mapped them to the protein family annotations (**Data S1**). We encoded each gene-pair adjacency and their orientations as a discrete character and reconstructed ancestral states at all internal nodes of the species tree with PastML^20^ (ML method = MPPA, character evolution model = EFT). This yielded, for each ancestral node of the species tree, a set of inferred adjacencies and the orientation of each pair. Each inference is assigned a posterior probability support. We converted the PastML-inferred adjacencies into an ancestral adjacency graph (nodes: ancestral genes; edges: adjacencies annotated with orientation and ≥0.5 posterior probability support). We linearized the graph using the AGORA strategy^26^: iteratively remove the lowest-support edges until contigs are oriented and non-branching. This approach yielded an initial set of chromosome-scale scaffolds.

PastML was also used to infer the ancestral karyotypes using the extant karyotypes as inputs. At each bifurcation of the species tree, we compared the inferred karyotype, the extant karyotypes of the daughters, and the reconstructed chromosome-scale scaffolds. When the reconstructed scaffolds mismatched the inferred karyotype, we manually adjusted the former by mapping them to the daughters’ chromosomes. We limited these reconstructions to N11, because chromosome-scale assemblies are unavailable for deeper nodes. The resulting chromosome-scale reconstructed ancestral genomes are available as **Data S3**.

**Genetic mechanisms underlying genome tramlining**

Using our protein family annotations (**Data S1**), we first established synteny blocks, *i.e.*, regions of conserved gene order, between the ancestral and daughter genomes at each bifurcation of the species tree. Next, we used in-house Python scripts to detect various genetic events.

**Chromosome fusion:** Two ancestral synteny blocks, which resided on separate chromosomes, are now found contiguously on a single chromosome in the daughter genome.

**Chromosome fission:** A contiguous ancestral synteny block is split across two different chromosomes in the daughter genome.

**Segmental inversion:** A contiguous block of multiple 1:1 orthologs has the opposite orientation in the ancestor and the daughter genomes.

**Paralog loss:** The ancestor had two paralogs (A1 and A2), but the daughter only retains A1.

**Gene duplication:** An ancestral gene corresponds to two or more paralogous genes in the daughter genome.

**Gene gain:** A gene is present in the daughter genome but has no ortholog in the ancestor. Of note, our analysis cannot distinguish *de novo* origin from horizontal transfers.

**Gene loss:** A gene present in the ancestor has no ortholog in the daughter genome.

**Gene inversion:** A 1:1 ortholog is found in the same synteny block, but its orientation is flipped in the daughter compared to the ancestor.

**Gene relocation:** A 1:1 ortholog is present in both the daughter and the ancestor, but its location on the chromosome differs from the flanking genes in the synteny block.

**Gene duplication followed by inversion:** An ancestral gene corresponds to two or more paralogous genes in the daughter genome. One paralog retains the ancestral synteny and orientation, the other is found in a new location in reverse orientation.

**Gene relocation followed by inversion:** A 1:1 ortholog is present in both the daughter and the ancestor, but its location on the chromosome differs from the flanking genes in the synteny block, and also its orientation is reversed.

**Mapping protein complexes and metabolic pathways**

We compiled a comprehensive dataset of characterized protein complexes, annotated for subunit compositions from the Complex Portal database^27^. This dataset included 2,420 complexes in human, 752 in mouse, 631 in yeast, 269 in fruit fly, 225 in thale cress, and 144 in roundworm. For each complex, the amino acid sequences of its subunits were obtained from UniProt^28^. We used reciprocal BLAST^7^ to map these subunits to our protein family annotations (≤1E−10 e-value with >50% of family members). Similarly, we obtained the enzyme composition data for characterized eukaryotic metabolic pathways from MetaCyc^29^ and similarly mapped them to our protein family annotations.

**Computing rigid body entanglement**

To quantify the core versus peripheral positioning of a subunit within the 3D structure of a complex, we counted the number of ways in which it can dissociate from the rest of the complex as a rigid body without causing steric clashes^30^. To that end, we kept the rest of the complex static, whereas the subunit of interest was translated by 4 Å in all directions around its center of mass. One thousand directions were uniformly sampled using the Fibonacci sphere algorithm^31^. For each iterative translation, we assessed steric clashes using backbone atoms. Clashes were defined as two atoms closer than the sum of their van der Waals radii. Any such shift causing ＞1% of all backbone atoms to clash was assigned as a forbidden dissociation (FD). The fraction of FDs is termed here Rigid Body Entanglement (RBE) and is a measure of subunit centrality. Indeed, for a subunit embedded in the complex core, most of the 1000 attempted separation modes will be forbidden.

**Supplementary Tables**

**Table S1.** Listed are 47 organisms with fully sequenced genomes included in the phylogenetic analysis, along with the database source, genome annotation number, genome size in base pairs, and the number of annotated proteins in the genome. BUSCO scores highlight genome completeness assessments.

| **Species** | **Source** | **Annotation number** | **BUSCO** |
| --- | --- | --- | --- |
| *Carpediemonas membranifera* | GenBank | GCA_019828565.1 | 89.0 |
| *Kipferlia bialata* | GenBank | GCA_003568945.1 | NA |
| *Paratrimastix pyriformis* | GenBank | GCA_027972325.1 | 76.3 |
| *Monocercomonoides exilis* | GenBank | GCA_001643675.2 | 75.3 |
| *Blattamonas nauphoetae* | GenBank | GCA_033064195.1 | 76.6 |
| *Naegleria gruberi* | GenBank | GCA_000004985.1 | 82.8 |
| *Naegleria lovaniensis* | GenBank | GCA_003324165.2 | 83.2 |
| *Diplonema papillatum* | GenBank | GCA_030068145.1 | 89.0 |
| *Perkinsela sp.* | GenBank | GCA_001235845.1 | 82.0 |
| *Bodo saltans* | TriTrypDB | GCA_001460835.1 | 86.9 |
| *Paratrypanosoma confusum* | TriTrypDB | GCA_002921335.1 | 96.2 |
| *Trypanosoma brucei* | TriTrypDB | GCA_000002445.1 | 99.2 |
| *Trypanosoma congolense* | TriTrypDB | GCA_000227395.2 | 93.8 |
| *Trypanosoma cruzi* | TriTrypDB | GCA_015033625.1 | 98.5 |
| *Trypanosoma equiperdum* | TriTrypDB | GCA_000300495.1 | 94.6 |
| *Trypanosoma evansi* | TriTrypDB | GCA_917563935.1 | 95.4 |
| *Trypanosoma grayi* | TriTrypDB | GCA_000691245.1 | 96.2 |
| *Trypanosoma melophagium* | TriTrypDB | GCA_022059095.1 | 100.0 |
| *Trypanosoma rangeli* | TriTrypDB | GCA_000492115.1 | 94.6 |
| *Trypanosoma theileri* | TriTrypDB | GCA_002087225.1 | 99.2 |
| *Trypanosoma vivax* | TriTrypDB | GCA_000227375.1 | 94.6 |
| *Blechomonas ayalai* | TriTrypDB | GCA_020509355.1 | 99.2 |
| *Phytomonas sp.* | GenBank | GCA_000582765.1 | 94.1 |
| *Strigomonas culicis* | GenBank | GCA_000442495.1 | 93.8 |
| *Angomonas deanei* | Ensembl | GCA_000442575.2 | 94.6 |
| *Crithidia fasciculata* | TriTrypDB | GCA_000331325.2 | 96.9 |
| *Leptomonas pyrrhocoris* | TriTrypDB | GCA_001293395.1 | 100.0 |
| *Leptomonas seymouri* | TriTrypDB | GCA_001299535.1 | 99.2 |
| *Endotrypanum monterogeii* | TriTrypDB | GCA_000333855.2 | 97.7 |
| *Porcisia hertigi* | TriTrypDB | GCA_017918235.1 | 98.5 |
| *Leishmania arabica* | TriTrypDB | GCA_000410695.2 | 98.5 |
| *Leishmania braziliensis* | TriTrypDB | GCA_000340355.2 | 98.5 |
| *Leishmania aethiopica* | TriTrypDB | GCA_000444285.2 | 95.4 |
| *Leishmania amazonensis* | TriTrypDB | GCA_000438535.1 | 97.7 |
| *Leishmania donovani* | TriTrypDB | GCA_003719575.1 | 100.0 |
| *Leishmania enriettii* | TriTrypDB | GCA_000410755.2 | 98.5 |
| *Leishmania gerbilli* | TriTrypDB | GCA_000443025.1 | 100.0 |
| *Leishmania infantum* | TriTrypDB | GCA_900500625.2 | 100.0 |
| *Leishmania major* | TriTrypDB | GCA_000002725.2 | 100.0 |
| *Leishmania martiniquensis* | TriTrypDB | GCA_000409445.2 | 100.0 |
| *Leishmania mexicana* | TriTrypDB | GCA_000234665.4 | 99.2 |
| *Leishmania orientalis* | TriTrypDB | GCA_017916335.1 | 99.2 |
| *Leishmania panamensis* | TriTrypDB | GCA_000755165.1 | 100.0 |
| *Leishmania tarentolae* | GenBank | GCA_009770625.1 | 99.2 |
| *Leishmania tropica* | TriTrypDB | GCA_000410715.1 | 100.0 |
| *Leishmania turanica* | TriTrypDB | GCA_000441995.1 | 100.0 |
| *Leishmania sp. Namibia* | GenBank | GCA_017918225.1 | 99.7 |

**Supplementary Figures**

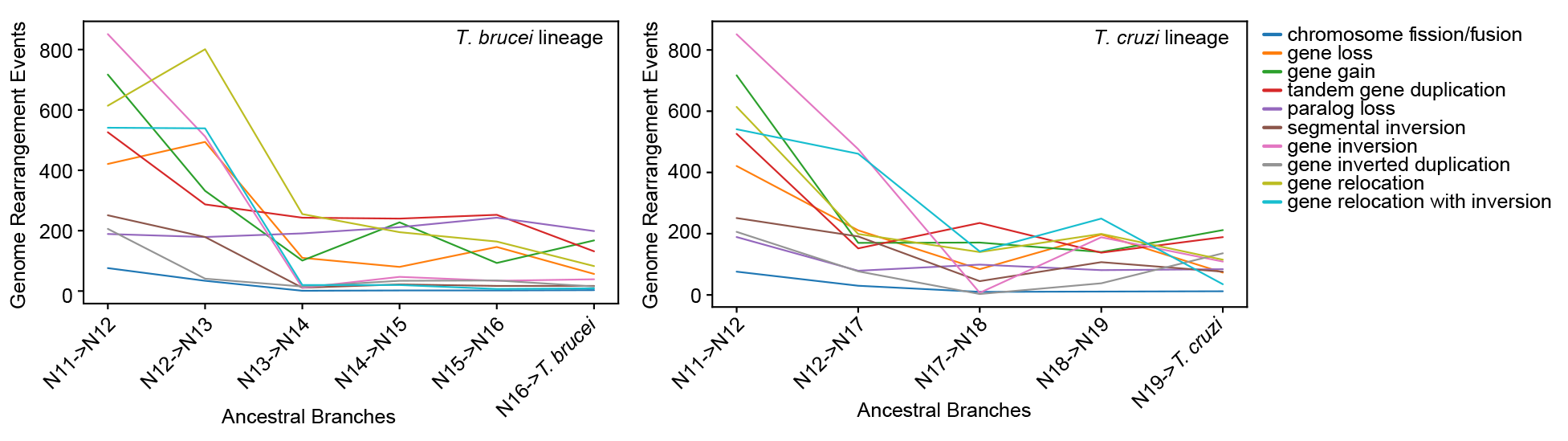

**Figure S1.** Lines represent the frequency of different phylogenetically reconstructed genome rearrangement events occurring at the ancestral branches of the Trypanosomatid lineage leading to *Trypanosoma brucei* (left) and *T. cruzi* (right). Plot features are the same as Figure 4F.

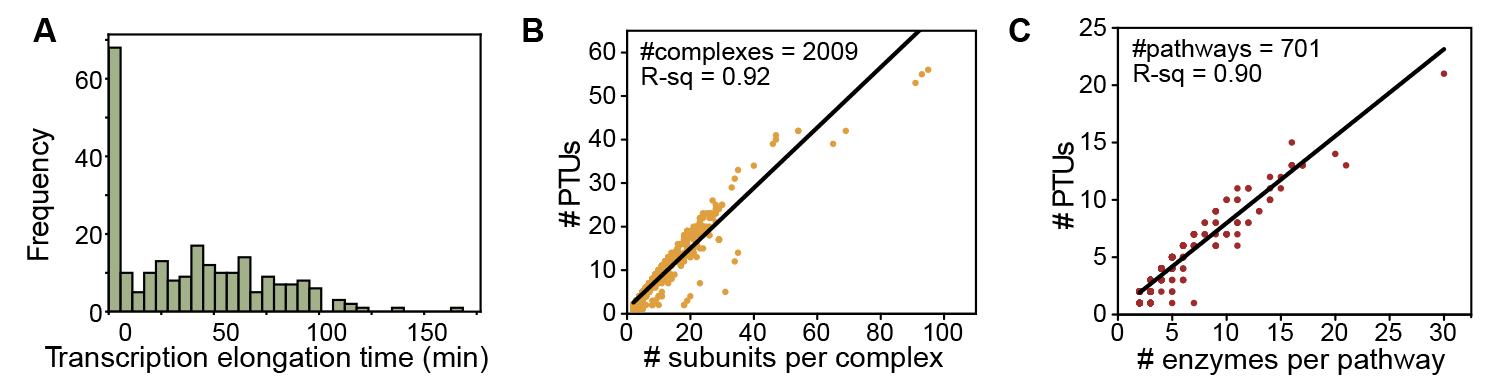

**Figure S2.** (**A**) A histogram plot depicts the distribution of transcription elongation times across *L. major* PTUs, assuming 60 nt/s transcription elongation speed. (**B-C**) A scatter plot shows the number of subunits per complex (**B**) and the number of enzymes per pathway (**C**) plotted against the number of PTUs in which their genes reside.
